# Nelfinavir induces cytotoxicity towards high-grade serous ovarian cancer cells, involving induction of the unfolded protein response, modulation of protein synthesis, DNA damage, lysosomal impairment, and potentiation of toxicity caused by proteasome inhibition

**DOI:** 10.1101/2021.10.25.465799

**Authors:** Mahbuba R Subeha, Alicia A Goyeneche, Prisca Bustamante, Michael A. Lisio, Julia V. Burnier, Carlos M. Telleria

**Author notes:** Correspondence should be addressed to CM Telleria.

## Abstract

High-grade serous ovarian cancer (HGSOC) is a significant cause of mortality among women worldwide. Traditional treatment consists of platinum-based therapy; however, rapid development of platinum resistance contributes to lower life expectancy, warranting newer therapies to supplement the current platinum-based protocol. Repurposing market-available drugs as cancer therapeutics is a cost- and time-effective way to avail new therapies to drug-resistant patients. The anti-HIV agent nelfinavir (NFV) has shown promising toxicity against various cancers; however, its role against HGSOC is unknown. Here, we studied the effect of NFV against HGSOC cells obtained from patients along disease progression and carrying different sensitivities to platinum. NFV triggered, independently of platinum sensitivity, a dose-dependent reduction of HGSOC cell number and viability, and a parallel increase in hypo-diploid DNA content. Moreover, a dose-dependent reduction of clonogenic survival of cells escaping the acute toxicity was indicative of long-term residual damage. In addition, dose- and time-dependent phosphorylation of H2AX indicated NFV-mediated DNA damage, which was associated with decreased survival and proliferation signals driven by the AKT and ERK pathways. NFV also mediated a dose-dependent increase in endoplasmic reticulum stress-related molecules associated with long-term inhibition of protein synthesis and concurrent cell death; such events were accompanied by a proapoptotic environment, signaled by increased phospho- eIF2α, ATF4, and CHOP, increased Bax/Bcl-2 ratio, and cleaved executer caspase-7. Finally, we show that NFV potentiates the short-term cell cycle arrest and long-term toxicity caused by the proteasome inhibitor bortezomib. Overall, our *in vitro* study demonstrates that NFV can therapeutically target HGSOC cells of differential platinum sensitivities via several mechanisms, suggesting its prospective repurposing benefit considering its good safety profile.

## Introduction

Ovarian cancer remains a significant source of morbidity and mortality, being the seventh most common cancer among women and the eighth leading cause of gynecologic-cancer-related death in the world [1]. Despite increasing advancement in diagnostics and therapeutic avenues, platinum-based therapy following tumor-debulking surgery persists as the backbone of ovarian cancer treatment for over fifty years. Although 70% of patients respond favourably to the initial platinum-based therapy, flaring of microscopic residual disease and emergence of platinum-resistance are inevitable within 18-24 months, which has contributed to an unimproved 5-year survival rate (47%) since the 1980s [2–5]. A maintenance therapy during the window of time between primary standard of care and the relapse of the disease may lead to improved patient survival.

Multiple efforts have been in place to develop consolidation and maintenance therapies for ovarian cancer, among which a few have been adopted into actual clinical practice, including poly (ADP-ribose) polymerase (PARP) inhibitors for BRCA1/2 mutant patients, and antiangiogenic agents for advanced-stage patients [6]. Epithelial ovarian cancer and its most lethal and prevalent subtype–high-grade serous ovarian cancer (HGSOC)–are considered heterogeneous diseases phenotypically and genetically [1, 7]. As such, a one-size-fits-all pharmacologic strategy may not be the optimal solution for every patient suffering from HGSOC. Thus, continual efforts need to be in place to upgrade the chemotherapeutic repertoire of ovarian cancer to offer multiple therapeutic options to patients at risk of evolving into platinum-resistant disease.

Novel drug development requires a decade-long time commitment and high-risk financial investments. Repurposing market-available drugs for newer indications can be an alternative and cost-effective strategy to fast track the availability of newer therapeutic options to drug-resistant as well as financially underprivileged patients [8]. An old drug with high repurposing potential is nelfinavir (NFV), which is a prototypical drug of the human-immunodeficiency virus protease inhibitor (HIV-PI) group designed to inhibit the requisite cleavage of HIV aspartyl protease– rendering the virions immature and non-infective [9]. NFV has been safely and extensively used as an oral anti-infective agent to treat acquired immunodeficiency syndrome (AIDS) in adult and pediatric patients since its approval in 1997 [10]. We have recently compiled *in vitro* and *in vivo* evidence supporting the efficacy of NFV against different cancers [11]. Among ten market available HIV-PIs, NFV has demonstrated the maximal anti-neoplastic efficiency via off-target effects as reported in disparate cancers such as lung [12], breast [13], prostate [14], multiple myeloma [15], leukemia [16], melanoma [17], and ovarian [18].

At the molecular level, the anti-cancer properties of NFV have been associated with multipronged mechanistic pathways among which upregulation of the stress of the endoplasmic reticulum (ER) and its associated unfolded protein response (UPR) has been a universal one. Depending on the type of cancer and treatment regimen, NFV has also been shown to modulate cell cycle, cell death, signal transduction pathways, autophagy, the proteasome pathway, the tumor microenvironment, oxidative stress, and multidrug efflux pumps [11].

In the current study, we report the preclinical efficacy of NFV against the most fatal histotype of ovarian cancer, high-grade serous (HGSOC). We define the effect of NFV against HGSOC cells of differential platinum sensitivities, patient origins, and disease progression. We further describe the mechanistic processes associated with NFV-induced toxicity against HGSOC cells, which involves DNA damage, reduced survival and proliferation signals, and a proapoptotic shift of the UPR associated to the ER stress pathway.

## Materials and Methods

### Cell culture and reagents

PEO1/PEO4/PEO6 and PEO14/PEO23, two sequentially obtained and spontaneously immortalized series of cell lines were utilized to explore the effects of NFV [19]. The original patients demonstrated different levels of disease progression and platinum sensitivities during the establishment of each cell line. The first patient was sensitive to cisplatin when the PEO1 cell line was established from ascitic fluid following 22 months of the last cisplatin-based therapy. Later, the patient was deemed clinically resistant to platinum while PEO4 and PEO6 cells were obtained respectively 10- and 3-months after the last cisplatin-based therapies. PEO14 cells were established from the ascitic fluid of a second patient during the chemo-naïve stage, and PEO23 cells were developed during the cisplatin-resistant stage of that patient 7 months after the last cycle of cisplatin-based therapy [20]. With written consent from Dr. Langdon (Edinburgh Cancer Research Center, Edinburgh, UK), PEO1, PEO4, and PEO6 cell lines were obtained from Dr. Taniguchi (Fred Hutchinson Cancer Center, University of Washington, Seattle, WA, USA). PEO14 and its longitudinally patient-matched pair PEO23 were obtained from Culture Collections, Public Health England (Porton Down, Salisbury, UK). All cells were cultured in RPMI-1640 (Mediatech, Manassas, VA, USA) supplemented with 5% fetal bovine serum (Atlanta Biologicals, Lawrenceville, GA, USA), 5% bovine serum (Life Technologies, Auckland, NZ), 1 mM sodium pyruvate (Corning, Corning, NY, USA), 2 mM L-Alanyl-L-Glutamine (Glutagro^TM^, Corning), 10 mM HEPES (Corning), 0.01 mg/mL human insulin (Roche, Indianapolis, IN, USA), 100 IU penicillin (Mediatech), and 100 μg/mL streptomycin (Mediatech). Cell culture was carried out at 37°C in a humidified incubator with 95% air/5% CO_2_ in standard adherent plastic plates. Autosomal short tandem repeat (STR) profiling markers were utilized for cell authentication, which demonstrated ≥ 80% match between the cell lines used in our study and the genotype of the original patients. The STR was carried out in the genetic core facility of the University of Arizona (Tucson, AZ, USA) [21]. The drugs used in this study were as follows: nelfinavir mesylate hydrate (NFV) (Sigma Chemical Co., St. Louis, MO, USA), cis-diamminedichloroplatinum II (cisplatin, Sigma), bortezomib (BZ) (Velcade®, Millennium Pharmaceuticals, Cambridge, MA, USA), tunicamycin (Sigma), puromycin dihydrochloride (Sigma), bafilomycin A1 (Cell Signaling Technology, Danvers, MA, USA), salubrinal (EMD Millipore, Billerica, MA, USA), and cycloheximide (Sigma). The drugs were dissolved in either dimethyl sulfoxide (DMSO, Sigma) or 0.9% sodium chloride solution (saline) (Sigma); the maximal concentration of DMSO in the cell culture was maintained at ≤ 0.1% (v/v).

### Cell proliferation and viability

Cell proliferation and viability were assessed via microcapillary cytometry. We described this methodology previously in detail [22]. Briefly, HGSOC cells were subjected to different treatments in triplicates or quadruplicates for different durations. After each treatment, cells were trypsinized and centrifuged to yield cellular pellets, which were washed and resuspended in phosphate-buffered saline (PBS), and mixed with the Muse™ Count & Viability Reagents (Luminex Corporation, Austin, TX, USA). The stained samples were analyzed with a microcapillary fluorescence cytometer (Guava® Muse® Cell Analyzer, Luminex), and the data were calculated by the Guava® Muse® Software (Luminex).

### Assessment of the sensitivity of the PEO cell line series to cisplatin

The sensitivity of the PEO cell lines to cisplatin was determined using a combination of short-term exposure of the cells to the drug followed by a long-term incubation of remaining live cells in cisplatin-free media. This assay allows determining the long-lasting toxic effects of cisplatin. The drug was stored in powder form until the time of treatment; it was then dissolved in saline at a concentration of 3333 μM. The drug was introduced into the media to reach final concentrations in the range of 1 to 50 μM. Saline was provided to vehicle-treated cells. Cells received cisplatin-infused media for 1 h, after which time media was removed, cells were washed with PBS, and media without cisplatin was provided for 72 h. The 1-h treatment time with cisplatin was selected to mimic the amount of time cisplatin is typically provided to a patient in a clinical setting. Thereafter, floating and adherent cells were collected and assayed for number and percent viability using fluorescence cytometry as explained above. Subsequently, 1000 viable cells for each treatment group were seeded in 6-well plates and cultured for 10-15 days until the number of cells/colony in the vehicle-treated plates was ≥ 50. At the end of the incubation period in cisplatin-free medium, the medium was aspirated, the cells were washed with PBS, and then fixed with 4% paraformaldehyde (PFA) for 30-45 minutes and stained with 0.5% (w/v) crystal violet (Sigma) for 20 minutes before being rinsed with tap water and dried at room temperature. Colonies having ≥ 50 cells were scored manually in an AmScope inverted light microscope with AmScope Software 3.7 (XD Series, United Scope LLC, Irvine, CA, USA) using 10x and 20x objectives. These values were input into the CalcuSyn software (Biosoft, Cambridge, UK) which calculated the half-maximal inhibitory concentration or IC_50_ (in µM) for each cell line. The average of two independent experiments performed in triplicate was used in the determination of the final IC_50_ value for each cell line (Table 1).

**Table 1.**
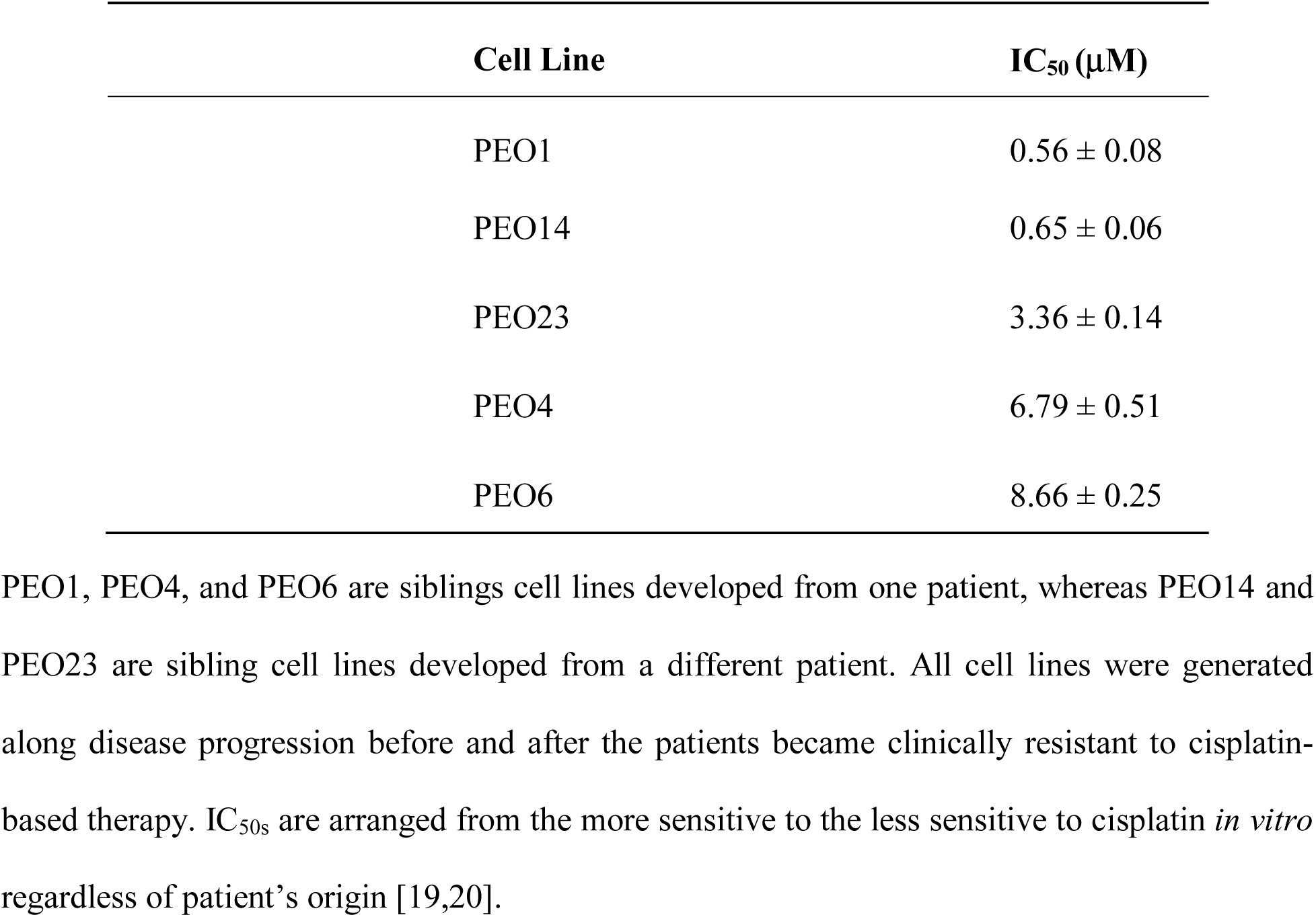
Concentration of cisplatin needed to achieve 50% reduction in clonogenic survival (IC_50_) of the HGSOC cell lines studied.

| Cell Line | $IC_{50}$ ($\mu M$) |
| --- | --- |
| PEO1 | $0.56 \pm 0.08$ |
| PEO14 | $0.65 \pm 0.06$ |
| PEO23 | $3.36 \pm 0.14$ |
| PEO4 | $6.79 \pm 0.51$ |
| PEO6 | $8.66 \pm 0.25$ |

PEO1, PEO4, and PEO6 are siblings cell lines developed from one patient, whereas PEO14 and PEO23 are sibling cell lines developed from a different patient. All cell lines were generated along disease progression before and after the patients became clinically resistant to cisplatin-based therapy. IC<sub>50s</sub> are arranged from the more sensitive to the less sensitive to cisplatin *in vitro* regardless of patient's origin [19,20].

### Cell cycle analysis

After NFV treatment, single cell suspensions were fixed with 4% PFA at room temperature for 1 h. Suspensions were centrifuged at 300 *g* for 5 minutes and pelleted cells were washed twice with PBS. A suspension containing 2 x 10^5^ cells was re-pelleted and resuspended in 0.2 mL of propidium iodide (PI) solution containing 7 U/mL RNase A, 0.05 mg/mL PI, 0.1 % v/v Triton X-100, and 3.8 mM sodium citrate (Sigma) for 20 min at room temperature or overnight at 4 °C protected from light. Cells were analyzed with the Guava® Muse® Cell Analyzer (Luminex), which takes advantage of the capacity of PI to stain DNA allowing detecting different DNA contents along the cell cycle. The cell cycle application of the Guava® Muse® software was used to analyze the results and to determine relative stages within the cell cycle. The PI-stained particles found containing hypo-diploid DNA content were considered to be in a Sub-G1 state likely representing apoptotic bodies.

### Clonogenic survival assay following NFV treatment

HGSOC cells were treated with increasing concentrations of NFV in triplicates or quadruplicates for 72 h; thereafter, the cells were trypsinized and counted using the method described in the cell proliferation and viability assay. For each treatment group, 1000 live cells were re-plated sparingly in 6-well plates in drug-free media and were incubated for 14-21 days to observe the colony-forming capacity of the cells withstanding the treatment. When the number of cells reached ≥ 50 per colony in the vehicle group, the experiment was terminated by discarding the media and fixing the cells with 4% PFA. The fixed cells were further stained using 0.5 % crystal violet, and the colonies were counted manually using an inverted light microscope as explained above.

### Measure of XBP1 mRNA splicing

RNA was isolated using RNAqueous™-4PCR DNA-free™ RNA Isolation for RT-PCR kit (Thermo Fischer Scientific, Waltham, MA, USA) following the manufacturer’s instructions. cDNA was synthetized using iScript (BioRad Laboratories, Inc, Hercules, USA) and quantified in a NanoDrop 2000 (Thermofisher). The forward primer for PCR amplification of spliced and total human Xbp1 mRNA was 5‘-CCTGGTTGCTGAAGAGGAGG-3’ and the reverse primer was 5’CCATGGGGAGTTCTGGAG-3’. For ACTB (β-actin), the primers were 5’ACAGAGCCTCGCCTTTG-3’ (forward) and 5’-CCTTGCACATGCCGGAG-3’ (reverse). The size of amplified unspliced Xbp1 mRNA is 145 base pairs (bp), the size of amplified spliced Xbp1 mRNA is 119 bp, and the size of amplified ACTB (β-Actin) is 110 bp. All primers were purchased from ID Technologies (Coralville, IA, USA). PCR reactions were performed in a 20 μL total volume reaction using SsoAdvanced Universal SYBR Green supermix (BioRad), 900 nM primer, 20 ng sample, and RT-PCR Grade Water (Invitrogen, Carlsbad, CA, USA). Using a C100 thermal cycler (BioRad), the following cycling profile was applied: 95°C for 3 min, followed by 35 cycles at 94°C for 30 s, 58°C (60°C for β-Actin) for 30 s, and 72°C for 30 s, with a final extension at 72°C for 5 min. No template control and no reverse transcriptase control were also included in each assay. The PCR products were visualized in 2% agarose gels, which were run at 100V and then imaged in a ChemiDoc MP (BioRad). The gels were stained with SYBR Safe DNA gel stain (Invitrogen). A Low DNA Mass Ladder (Invitrogen) was used to determine the size of the PCR products.

### Western blot analysis

Following treatment, HGSOC cells were washed with ice-cold PBS, scraped, collected, and centrifuged to yield pellets, which were stored in -80°C. Protein lysates were extracted from the pellets using NP40 lysis buffer, and 20 μg of proteins per sample were resolved in 10 or 12% gels (TGX™ FastCast™ Acrylamide kit, Bio-Rad) via electrophoresis. The resolved proteins were transferred to Immuno-Blot® PVDF membranes using a Trans-Blot® Turbo™ Transfer System (BioRad). Membranes were incubated at 4°C overnight in primary antibodies against GRP78, CHOP, IRE1α, PERK, p-eIF2α, eIF2α, ATF4, LC3II, Bcl-2, Bax, Caspase-7, p-AKT (Ser-473), p-AKT (Thr-308), AKT, p-ERK, ERK, ATF6, p27^kip1^, Cdk2, p-Cdk2, γH2AX, p-KAP1, puromycin, and β-actin. Thereafter, membranes were washed and re-incubated with secondary antibodies, and protein detection was performed via a ChemiDoc Imaging System (BioRad) using chemilluminescence (Clarity Western ECL Imaging System, BioRad) (see Table S1 for details about the origin and dilutions of the used antibodies). Ultraviolet activation of the TGX stain-free gels on a ChemiDoc MP Imaging System (BioRad) was used to control for proper loading In short, the pre-cast gels include unique trihalo compounds that allow rapid fluorescence detection of proteins without staining. The trihalo compounds react with tryptophan residues in a UV-induced reaction to produce fluorescence that is detected on the PVDF membranes [23]).

### Puromycin incorporation assay

The puromycin incorporation assay is a non-radioactive method of quantifying mRNA translation rate. Puromycin is an aminoacyl-tRNA mimetic that can occupy the site A of the ribosome during mRNA translation and thereby terminates the process prematurely. However, short-term exposure enables conjugation of puromycin with the nascent polypeptide chains generating short-lived puromycylated peptides that are released from the ribosome and can be detected by an anti-puromycin antibody on immunoblots. As one molecule of puromycin is incorporated into each released nascent polypeptide, puromycin incorporation is deemed as a sensitive indicator of ongoing mRNA translation rate [24, 25]. In our experiments, puromycin was added to the culture media at a final concentration of 1 μM at 37°C, 30 min prior to the termination of experiments. Thereafter, the cells were collected and processed for the detection of puromycylated proteins by immunoblot.

### Autophagic flux

We studied autophagic flux as previously described in our laboratory [26]. Briefly, autophagic flux assay is performed by tracking the expression of the autophagosomal membrane-associated protein LC3II in response to a drug treatment, with or without the presence of an inhibitor of the lysosomal function. An increase in the levels of LC3II in response to a drug treatment may indicate either increased autophagy induction or impaired autophagosome removal by the lysosomes. Hence, autophagic flux is a better measure of the autophagic process, as it determines LC3II turnover in the presence or absence of the lysosome inhibitor bafilomycin A1. If LC3II levels raise further in the presence of the studied drug plus bafilomycin A1 when compared to cells treated only with the experimental drug, this means that the rate of autophagy or autophagic flux is increased. In contrast, no change in the expression of LC3II during co-treatment with the lysosome inhibitor indicates accumulation of autophagosomes because of lysosome impairment caused by the drug under study. In our experiments, NFV-treated HGSOC cells were further exposed or not to 100 nM bafilomycin A1 for 1 h before the termination of the experiment. Thereafter, the cells were collected and processed for the detection of LC3II by immunoblot.

### Drug interaction analysis

To determine whether there is pharmacological interaction between NFV and BZ, a drug-interaction assay was performed on platinum-sensitive PEO1, and on the less-sensitive to platinum, PEO4 cells. Using total number of cells as a variable, data were analyzed through algorithms in the CalcuSyn software (Biosoft), which uses the combination index (CI) method for predicting drug interaction [27]. We previously described in detail the calculation of the CI [28]. In brief, for a specific drug association, a CI>1 is considered antagonistic, CI=0 means no drug interaction, CI=1 indicates additivism, whereas CI<1 denotes synergism.

### Statistical analysis

For tests involving western blot analysis, the experiments were repeated at least twice with similar outcome. Numerical data are expressed as the mean ± SEM. Differences were considered significant if P<0.05. GraphPad Prism 9 (Graphpad Software, La Jolla, CA, USA) allowed for statistical analysis of data using one-way ANOVA followed by Tukey’s multiple comparison test, or two-way ANOVA followed by Bonferroni’s multiple comparison test.

## Results

### Nelfinavir inhibits growth, reduces viability, increases hypo-diploid DNA content, and blocks clonogenic survival of high-grade serous ovarian cancer (HGSOC) cells regardless of platinum sensitivity

The overall cytotoxicity of NFV was assessed on HGSOC cells of varying cisplatin sensitivities, which are depicted in Table 1. Of the five tested cell lines, PEO1, PEO4, and PEO6 were derived from one patient, whereas PEO14 and PEO23 were derived from a second patient. PEO1 demonstrated to be the most sensitive to cisplatin, whereas PEO6 showed to be the least sensitive. Our results also indicate that the cell lines, established at different stages of disease progression, recapitulated in vitro the cisplatin sensitivity of the original patients during the time of procuring the cells from ascites, with cisplatin sensitivity of PEO1>PEO4>PEO6 for patient one, and PEO14>PEO23 for patient two.

In this study we assessed cell toxicity in a broad manner, including abrogation of their reproductive capacity (cytostasis), and the actual dying of the cells (lethality) which can be acute and visualized upon short-term incubation (within 72 h), or long-lasting irreversible reproductive impairment visualized in long-term incubations (as observed in clonogenic survival assays). Acute cytotoxicity of NFV towards HGSOC cells was investigated via cell proliferation and viability assays following 72 h of treatment. We observed that NFV decreased the total number of cells and the percent viability of all the cell lines in a concentration-dependent manner regardless of their platinum sensitivity (Fig. 1A and B). We further determined that higher concentrations of NVF reduce cellular viability in association with the accumulation of hypo-diploid DNA content (a.k.a. Sub-G1 DNA content), denoting that the cells are likely dying by apoptosis (Fig. 1C). Finally, the long-lasting reproductive impairment caused by NFV in HGSOC cells was reflected by the fact that cells that remained alive after 72-h exposure to NFV had reduced clonogenic capacity when incubated in NFV-free media for 15-21 days (Fig. 1D); the IC_50s_ of NFV reported for these clonogenic survival assays are very similar among the cell lines studied (10-16 μM) regardless of their different sensitivities to platinum (Table S2). In summary, NFV is toxic towards HGSOC cells regardless of their sensitivities to cisplatin involving short-term reduction in viability and leading to a long-lasting impairment of their reproductive capacities.

**Figure 1.**
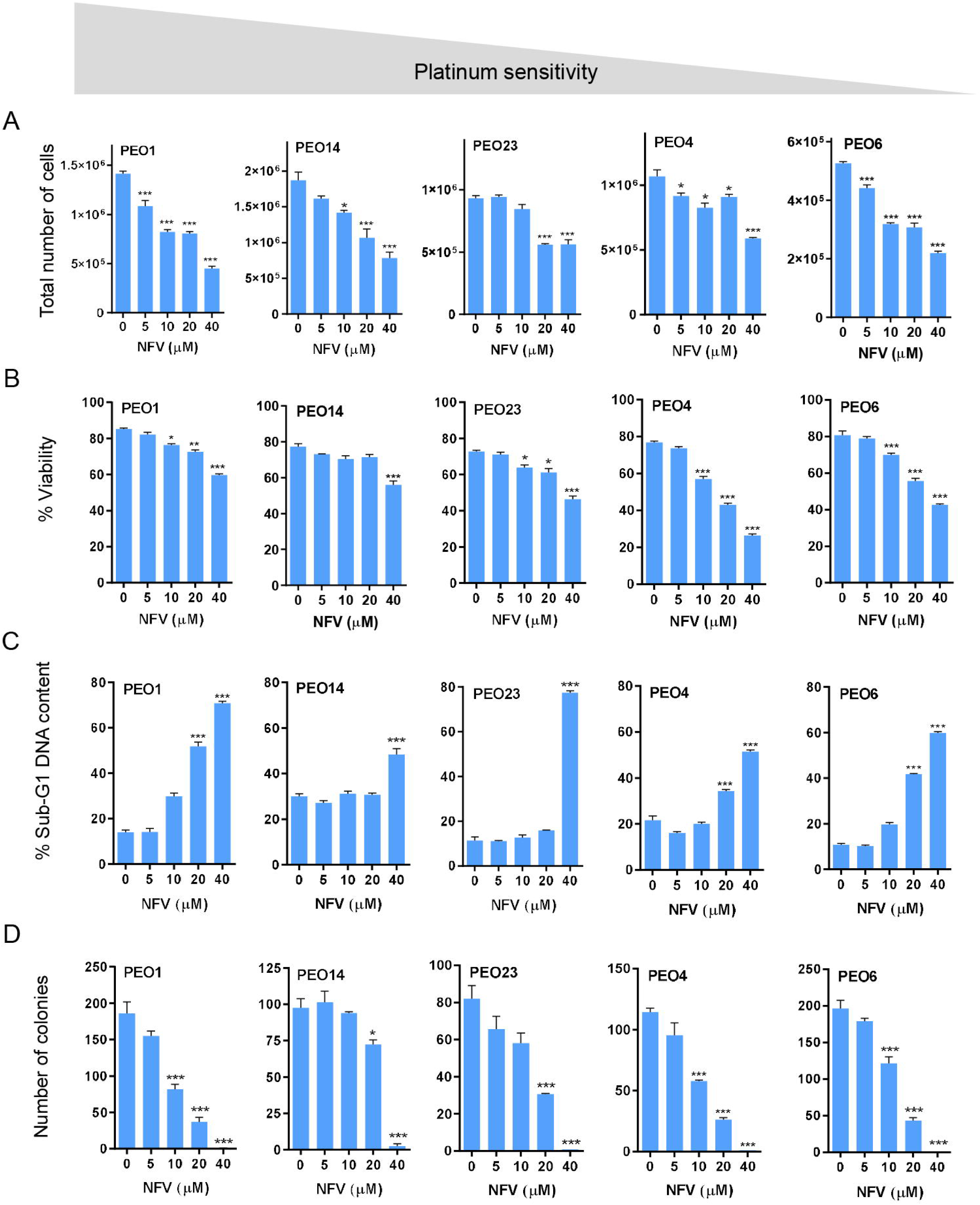
HGSOC cells of differential platinum sensitivities were plated in six-well plates in triplicate and, when exponentially growing, were subjected to treatment with the depicted concentrations of nelfinavir (NFV) for 72 h. At the end of the experiment, the cells were trypsinized and subjected to microcytometry analysis to attain total number of viable cells (A), and percent viability (B). A fraction of the cellular content collected at the end of the incubation with NFV was stained with a solution of propidium iodide (PI) and subjected to cell cycle analysis to determine hypodiploid DNA content (C). Finally, 1000 viable cells obtained at the end of the experiment were subjected to a clonogenic survival assay in the absence of treatment to determine delayed toxicity of NFV (D). *p<0.05; ***p<0.001 vs. control (One-way ANOVA followed by Tukey’s Multiple Comparison Test). The gray area that goes from higher to lower signifies that the cells are depicted in order of their decreasing sensitivity to the toxic effects of cisplatin as presented in Table 1.

### Nelfinavir induces cell cycle arrest, triggers endoplasmic reticulum (ER) stress and the unfolded protein response (UPR), and impairs the function of lysosomes without affecting autophagic flux

To determine whether the impaired cell proliferation induced by NFV is in the short-term associated with cell cycle arrest, we incubated HGSOC cells of different platinum sensitivities with increasing concentrations of NFV for 72 h. We measured the expression of the cyclin-dependent kinase (Cdk) inhibitor p27^kip1^ and found it increased in a concentration-dependent manner in all cell lines studied (Fig. 2A). Of interest, we found that the increase in p27^kip1^ associates with a decline in the phosphorylation levels of Cdk2 on Thr-160, which causes a reduction in the activity of Cdk2 [29], thus suggesting inhibition by NFV of the G1/S transition (Fig. S1). Moreover, we decided to explore whether NFV-associated cell growth inhibition involves the induction of the ER stress response because this pathway has been reported to be ubiquitously activated by NFV in multiple cancers [11]. We observed that, in all cell lines, NFV upregulates GRP78 (glucose-regulated protein, 78 kDa), which is a member of the family of heat shock proteins of 70 kDa, also termed heat shock 70 kDa protein 5 (HSPA5), and considered a master chaperone [30]; concomitantly, we detected NFV-induced upregulation of CCAAT-enhancer-binding protein homolog protein (CHOP) [31] (Fig.2A). Both GRP78 and CHOP are postulated to balance the stress of the ER in opposite manners, with CHOP being a pro-cell death factor and GRP78 a pro-survival factor [32]. This is consistent with the concept that ER stress is primarily a pro-survival mechanism, yet in excess it facilitates cell death [33].

**Figure 2.**
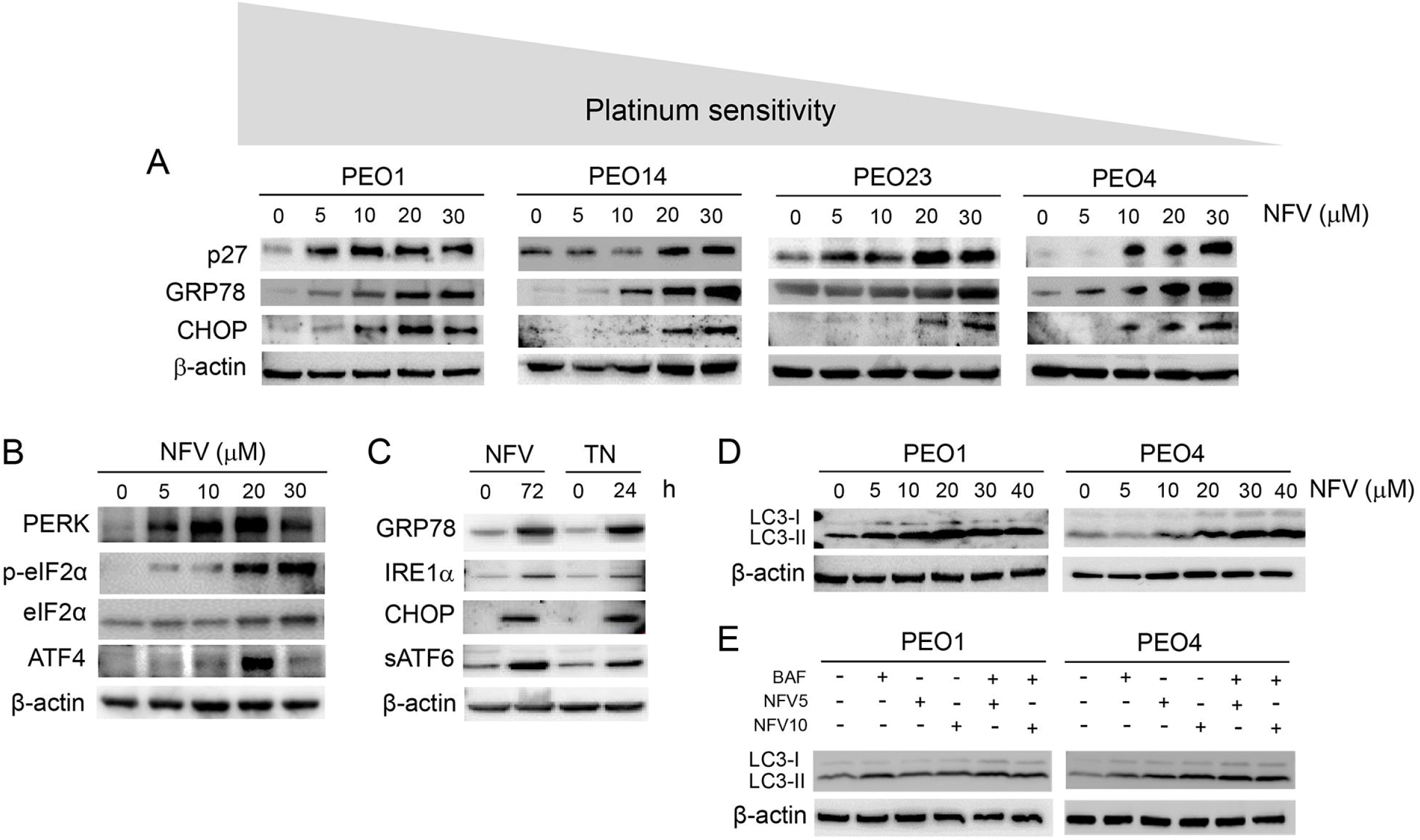
(A) PEO cells with different sensitivities to cisplatin were exposed to 5, 10, 20, or 30 µM nelfinavir (NFV) for 72 h. In all cell lines, NFV increased the expression of cell cycle inhibitor p27^kip1^, and ER stress related proteins GRP78, and CHOP. The gray area that goes from higher to lower signifies that the cells are depicted in order of their decreasing sensitivity to the toxic effects of cisplatin as shown in Table 1. (B) Expression of proteins involved in the ER response as studied in PEO1 cells exposed to various doses of NFV for 72 h. (C) Exposure of PEO1 cells to 20 μM NFV for 72 h increases ER stress-related proteins in a similar manner than 2 μg/mL of the known ER stressor tunicamycin (TN) does after 24 h exposure. (D) In PEO1 and PEO4 cells treated with various concentrations of NVF for 72 h, the autophagosome-related-protein LC3II increases in response to NFV in a concentration-dependent manner. (E) PEO1 and PEO4 cells were treated with 5 or 10 µM NFV for 72 h, in the absence or presence of 100 nM of the lysosome inhibitor bafilomycin A1 (BAF), which was added 1 h before the termination of the experiment. In both PEO1 and PEO4 cells, the induction of LC3II triggered by NFV was not augmented further by the presence of BAF. All results presented are representative of at least two independent experiments that had a similar outcome. NFV5, 5 μM NFV; NFV10, 10 μM NFV.

To maintain homeostasis when the load of unfolded proteins exceeds the folding capacity of the ER, GRP78 detaches from the ER membrane sensors PERK, IRE1, and ATF6, and activates the UPR [34–38]. These pathways are an adaptive response aimed at restoring homeostasis by inhibiting global protein synthesis, promoting enhanced expression of chaperone proteins, and favoring the degradation of misfolded proteins in the proteasome. We show that the PERK/eIF2α/ATF4 pathway of the UPR is stimulated by NFV in a concentration-dependent manner (Fig. 2B). We also compared members of the UPR in response to NFV against that caused by a recognised activator of the UPR, tunicamycin (TN), which is a glycosidase inhibitor causing accumulation of non-glycosylated proteins in the ER [39, 40]. We found that, similarly to TN, NVF increased GRP78, CHOP, and the other two pathways of the UPR, one involving the endonuclease IRE1α, and the other mediated by activation of ATF6 formed upon its trafficking from the ER to the Golgi apparatus where it is cleaved to release the soluble transcription factor (sATF6) (Fig. 2C).

Multiple studies have reported activation of autophagy in cancer cells in response to NFV treatment [12, 41]. Autophagy is an evolutionarily conserved biological mechanism aimed at disintegrating cellular organelles and bulky misfolded proteins, and recycling macromolecules to compensate for energy and nutrient deprivation [42, 43]. Furthermore, the PERK-eIF2α arm of ER stress has been associated with the modulation of autophagy [33]. To investigate if the ER stressor NFV affects autophagy in HGSOC cells, we treated platinum-sensitive PEO1 cells and its less sensitive patient-matched pair PEO4 with increasing concentrations of NFV for 72 h. We observed a concentration-dependent increase in the level of LC3II protein in both cell lines in response to NFV, suggesting accumulation of autophagosomes (Fig. 2D). We detected a similar increase in LC3II triggered by NFV in the other siblings HGSOC cell lines studied, PEO14 and PEO23 (Fig. S2). Increased level of LC3II however can indicate either a true increase in the dynamic process of autophagy (a.k.a. autophagic flux), or instead, an impairment of lysosomal activity [44]. To differentiate between these two outcomes, we performed an autophagic flux assay by co-treating the cells with NFV in the presence or absence of the lysosome inhibitor bafilomycin A1. NFV did not further enhance the level of LC3II when bafilomycin A1 was added to the treatment (Fig. 2E), suggesting that it likely impairs lysosomal function instead of enhancing autophagic flux.

### ER stress response induced by nelfinavir is associated with cleavage of executioner caspase-7 and increased pro-apoptotic Bcl-2 family member Bax in a time- and concentration-dependent manner

To further characterize the NFV-induced ER stress and associated UPR in HGSOC cells, we conducted a time-course experiment utilizing a single concentration of NFV. As predicted, NFV caused a time-dependent increase of ER stress-related proteins GRP78, IRE1α, ATF4, and CHOP, which was concomitant to the cleavage of executer caspase-7 and the increase in Bax/Bcl-2 ratio (Fig. 3A); this result is in agreement with the attributed pro-apoptotic function of ATF4 and CHOP during ER stress [37]. We also studied the expression of executer caspase-3, whose cleavage was not clearly detected (data not shown). We also found that the activation of caspase-7 and the increase in the Bax/Bcl-2 ratio by NFV is concentration dependent and occurs in both PEO1 and PEO4 sibling cells, which display highly different sensitivities to platinum (Table 1) (Fig. 3B). To further demonstrate the activation of the ER stress response as mediator of the overall toxicity displayed by NFV against HGSOC cells, we show that the reduction in viability induced by 20 μM NFV in PEO1 cells is abrogated by salubrinal, a known small molecule that protects cells from ER stress [45] (Fig. S3).

**Figure 3.**
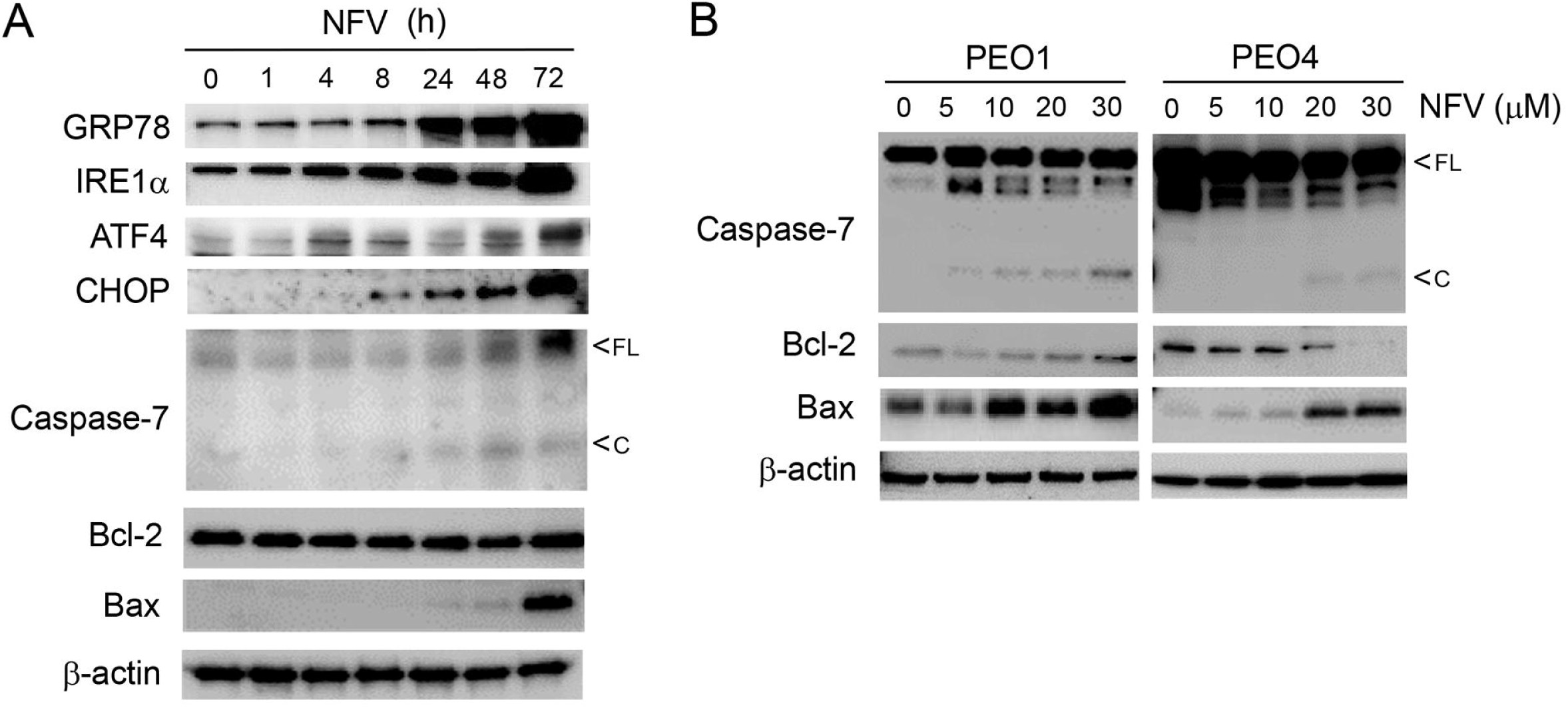
(A) PEO1 cells treated with 20 μM NFV for 72 h depict a time-dependent increase in ER stress related proteins GRP78, IRE1α, ATF4, and CHOP, while inducing the cleavage of executer caspase-7, and an increase in the Bax/Bcl-2 ratio. (B) NFV treatment of both PEO1 and PEO4 cells for 72 h cause a dose-dependent increase in cleaved caspase-7 while increasing the Bax/Bcl-2 ratio. All results presented are representative of at least two independent experiments that had a similar outcome.

### Nelfinavir toxicity is associated with short-term sustained mRNA translation that contributes to the UPR, followed by long-term concentration-dependent inhibition of global protein synthesis

One of the primary aims of the UPR is to reduce further protein load in the ER by shutting down global protein synthesis yet resume selective cap-independent translation to facilitate cellular recovery from the ongoing proteotoxic stress [33, 37]. To understand the effect of NFV on the dynamics of protein synthesis, NFV-treated PEO1 cells were subjected to a puromycin incorporation assay to assess mRNA translation. NFV inhibited mRNA translation in a concentration- (Fig. 4A) and time-(Fig. 4B) dependent manner. However, protein synthesis still was sustained during the first 4 h of NFV treatment, yet declined thereafter because puromycin incorporation after 24 h (Fig.4B) was similarly low as that observed when puromycylation was abrogated by the presence of the protein synthesis inhibitor cycloheximide (CHX) during the first 4 h of NVF treatment (Fig. 4C).

**Figure 4.**
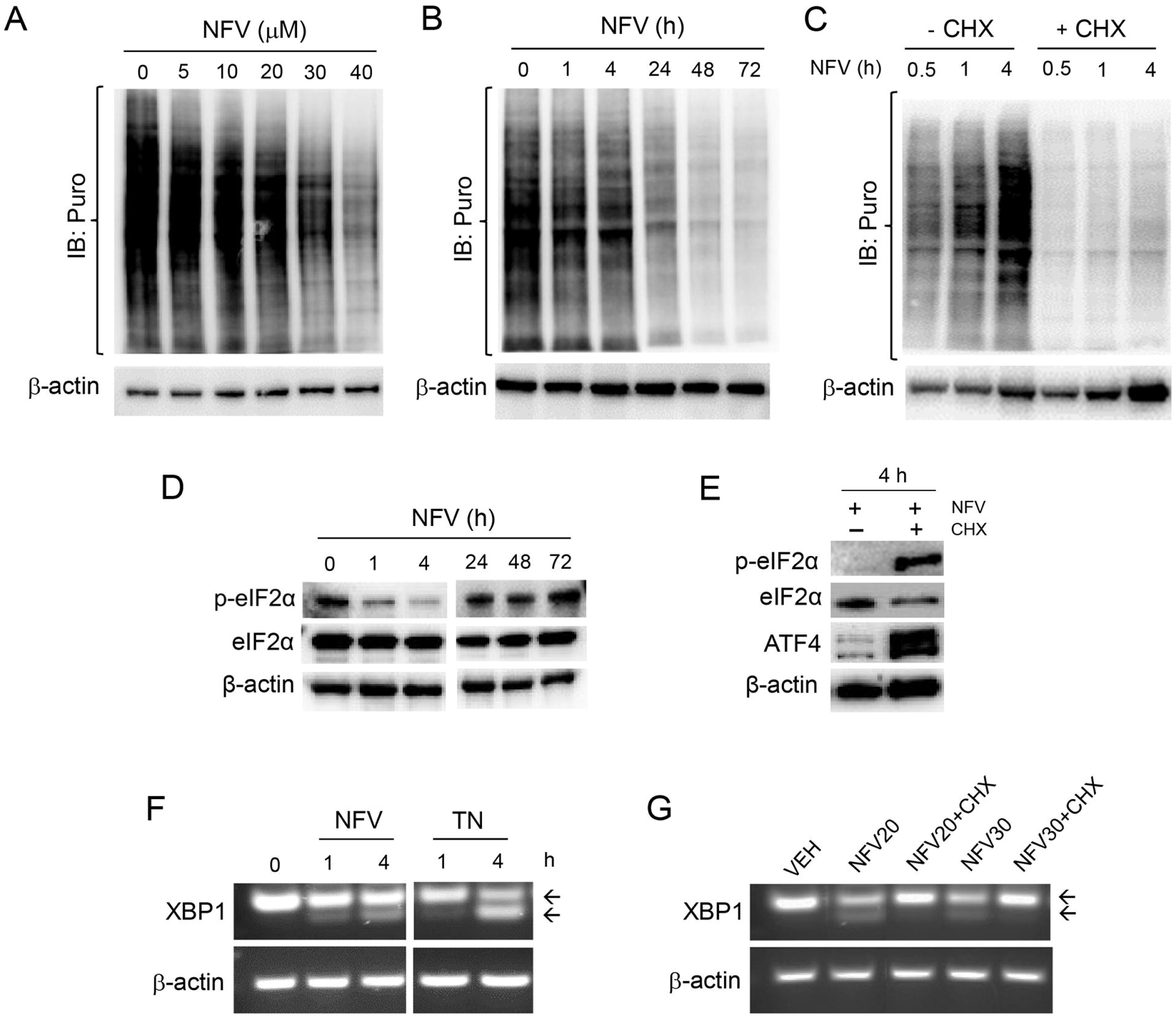
NFV, in a concentration- (A) and time-dependent manner (B), triggers a decrease in the incorporation of puromycin (Puro) into newly synthesized peptides. (C) The incorporation of Puro is sustained in the presence of 20 μM NFV during the first 4 h of treatment, but it is abrogated by the simultaneous presence 10 μg/mL of the protein synthesis inhibitor cycloheximide (CHX). (D) Effect of 20 μM NFV on the phosphorylation of eIF2α on Serine 51 (p-eIF2α) along a 72 h incubation period. (E) A short-term (4 h) treatment with 20 μM NFV associates with low expression of p-eIF2α and downstream transcription factor ATF4, yet both p-eIF2α and ATF4 radically increase with the simultaneous presence of CHX (10 μg/mL). (F) Effect of 20 μM NFV on the splicing of XBP1 mRNA assessed by RT-PCR; TN, tunicamycin (2 μg/mL). Arrows indicate total and spliced XBP1 mRNA variants. (G) Splicing of XPB1 mRNA in cells co-incubated for 4 h with NFV and CHX (10 μg/mL); NFV20, 20 μM; NFV30, 30 μM. All results presented were performed in PEO1 cells and are representative of two experiments that had a similar outcome.

Reduction of global protein synthesis upon ER stress induction occurs because of the phosphorylation of eIF2α. This is a polypeptide chain translation initiator factor that limits protein synthesis under conditions of cellular stress based on its capacity to become phosphorylated on serine 51 [46], thus limiting the availability of eIF2α needed for translation initiation [47]. The basal levels of eIF2α phosphorylation on serine 51 (p-eIF2α) are elevated, but were rapidly, yet temporarily diminished by NFV for about 4 h without affecting total eIF2α levels (Fig 4D); this was concurrent with the sustained incorporation of puromycin (Fig. 4C). This sustained protein synthesis at the beginning of the treatment with NFV was confirmed by reduced puromycin incorporation after 4 h of treatment with NFV (Fig 4C) and the sharp increase in p-eIF2α and downstream transcription factor ATF4 in the presence of CHX (Fig. 4E).

Despite in the long-term NFV-induced ER stress leads to a decline in global protein synthesis (Fig. 4B), we asked the question as to whether the sustained levels of protein synthesis during the first 4 h following NFV treatment, corroborated by reduced p-eIF2α and sustained puromycin incorporation, further induces ER stress. We answered this question by exposing PEO1 cells to NFV for 4 h, and measured a non-translatable readout of ER stress, the total, and spliced mRNA coding form XPB1, in the presence or absence of CHX. RT-PCR revealed NFV- mediated early splicing of XBP1 mRNA, the downstream target of IRE1α, which was similar to early XBP1 mRNA splicing induced by the known ER stressor tunicamycin (TN) (Fig. 4F). This cleavage, however, was prevented by the presence of CHX (Fig. 4G), suggesting that proteins accumulated during the first 4 h of NFV treatment participate at least in part in the causation of ER stress and the unleashing of the UPR. Taken together, these results provide evidence for a cross talk between ER stress and the modulation of protein synthesis dynamics in HGSOC cells in response to NFV.

### Nelfinavir inhibits AKT and ERK phosphorylation and triggers DNA damage

Elevation of cell survival and proliferation are typically favoured by an upregulation of AKT and ERK signaling pathways. NFV has been shown to reduce the phosphorylation of AKT and ERK in various cancers [13, 48]. In this study, we observed a concentration-dependent reduction of phosphorylation of both AKT and ERK in the siblings PEO1 and PEO4 cell lines carrying different sensitivities to cisplatin (Fig. 5). The decline in the activation of AKT (suggested by the dephosphorylation of its two activation sites, Ser-473 and Thr-308), and ERK (also undergoing dephosphorylation) was associated with NFV-mediated DNA damage response in both PEO1 and PEO4 cells, evidenced by a concentration-dependent increase of Ser-139 phosphorylated H2AX (γH2AX) and Thr-824 phosphorylated KAP1 (Fig. 5A). Induction of DNA damage in PEO4 cells is significant as this cell line has a restoration of a DNA-damage repair mechanism [49], which is inherently deficient in its patient-matched pair PEO1, thus conferring the reduced sensitivity to cisplatin observed in PEO4 cells (Table 1). We further demonstrated that the induction of DNA damage by NFV is time dependent (Fig. 5B).

**Figure 5.**
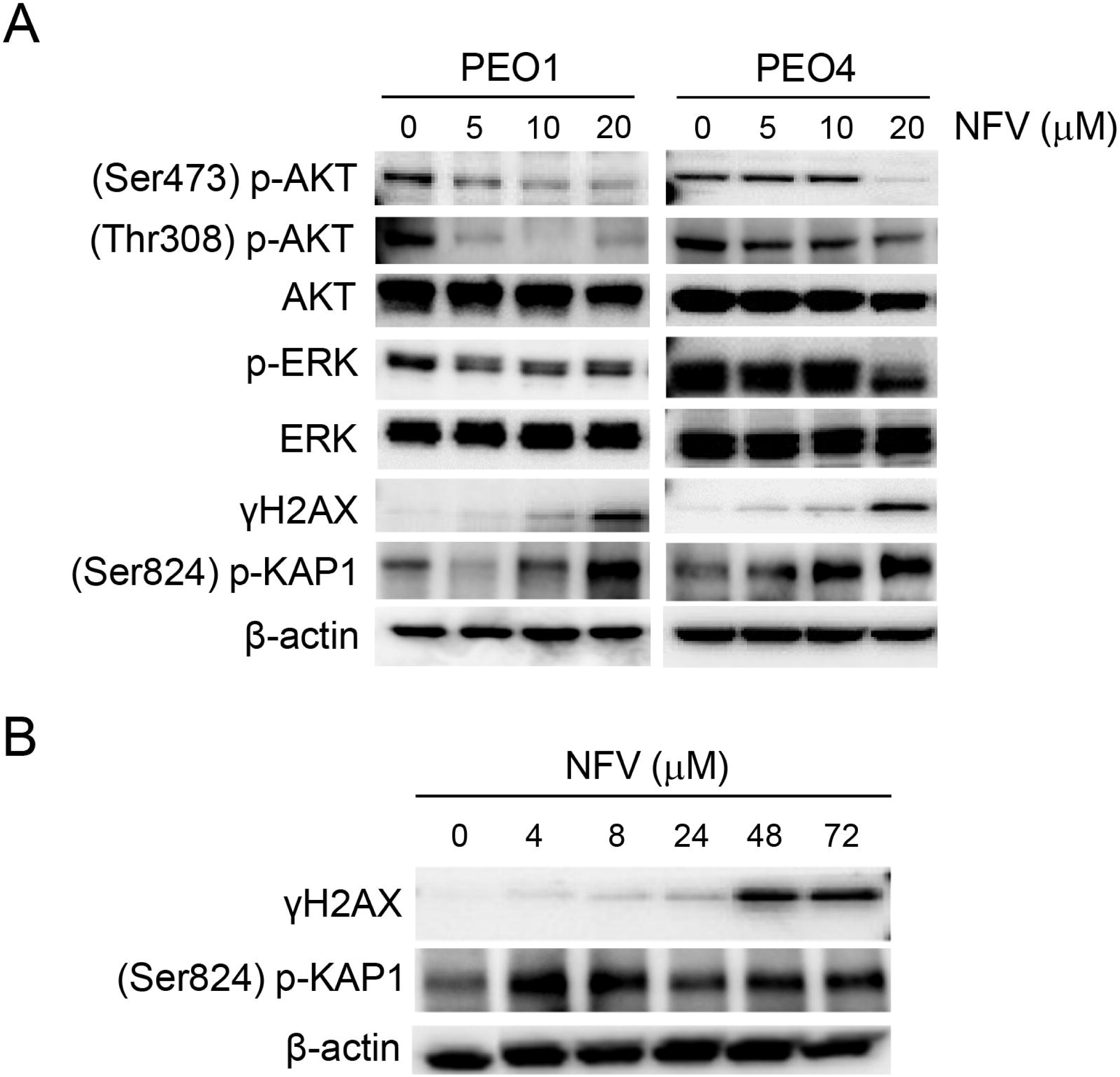
(A) PEO1 and PEO4 cells were incubated with the depicted concentrations of NFV for 72 h. Protein expression studied by western blot demonstrates concentration-dependent decrease in the phosphorylation of AKT and ERK in both sibling cell lines, which was accompanied by a concurrent increase of DNA damage markers γH2AX and (Ser824) p-KAP1. (B) Time course showing the induction of γH2AX and (Ser824) p-KAP1 in PEO1 cells treated with 20 μM NFV. All results presented are representative of at least two independent experiments that had a similar outcome.

### Nelfinavir potentiates the toxicity of the proteasome inhibitor bortezomib without modifying its proteasome inhibitory capacity

During the pro-survival phase, ER stress is relieved by activating ER-associated degradation (ERAD), whereby misfolded proteins are ubiquitinylated and translocated to the 26S proteasomes to undergo protein degradation thus contributing to a reduction in protein overload [33]. Previous studies have shown that the blocking of the 26S proteasomes may further increase the level of misfolded proteins and drive ER stress elicited by ER stressors from a pro-survival phase toward a proapoptotic one [26]. Likewise, we rationalized that NFV, acting as an ER stressor, could potentiate the toxicity of the proteasome inhibitor bortezomib (BZ).

PEO1 and PEO4 cells were exposed to combination treatments of varying concentrations of NFV and BZ used in a fixed ratio (1000:1) to assess drug interactions via the combination index of Chou and Talalay by utilizing the total cell count as a readout [27]. Drug synergism was predicted in PEO1 cells at a combination of 10 μM of NFV and 10 nM of BZ (CI=0.72). Similarly, drug synergism was predicted in PEO4 cells at a combination of 5 μM of NFV and 5 nM of BZ (CI=0.56). In both experiments, the concentrations of NFV that synergized with BZ where within the concentrations anticipated to be achieved in the clinic [41]. Cell proliferation assay demonstrated potentiation of toxicity with the predicted combinations of NFV and BZ in PEO1 (Fig. 6A) and PEO4 cells (Fig. 6E). By studying cell cycle distribution, we found that in both cell lines the combination of NFV/BZ caused accumulation of cells in the G1 phase of the cell cycle (Fig. 6B and F). This G1 cell cycle arrest caused by the drug combination was associated with the accumulation of the cell cycle inhibitor p27^kip1^ and the DNA damage markers γH2AX and p-KAP1 (Fig. 6C and G).

**Figure 6.**
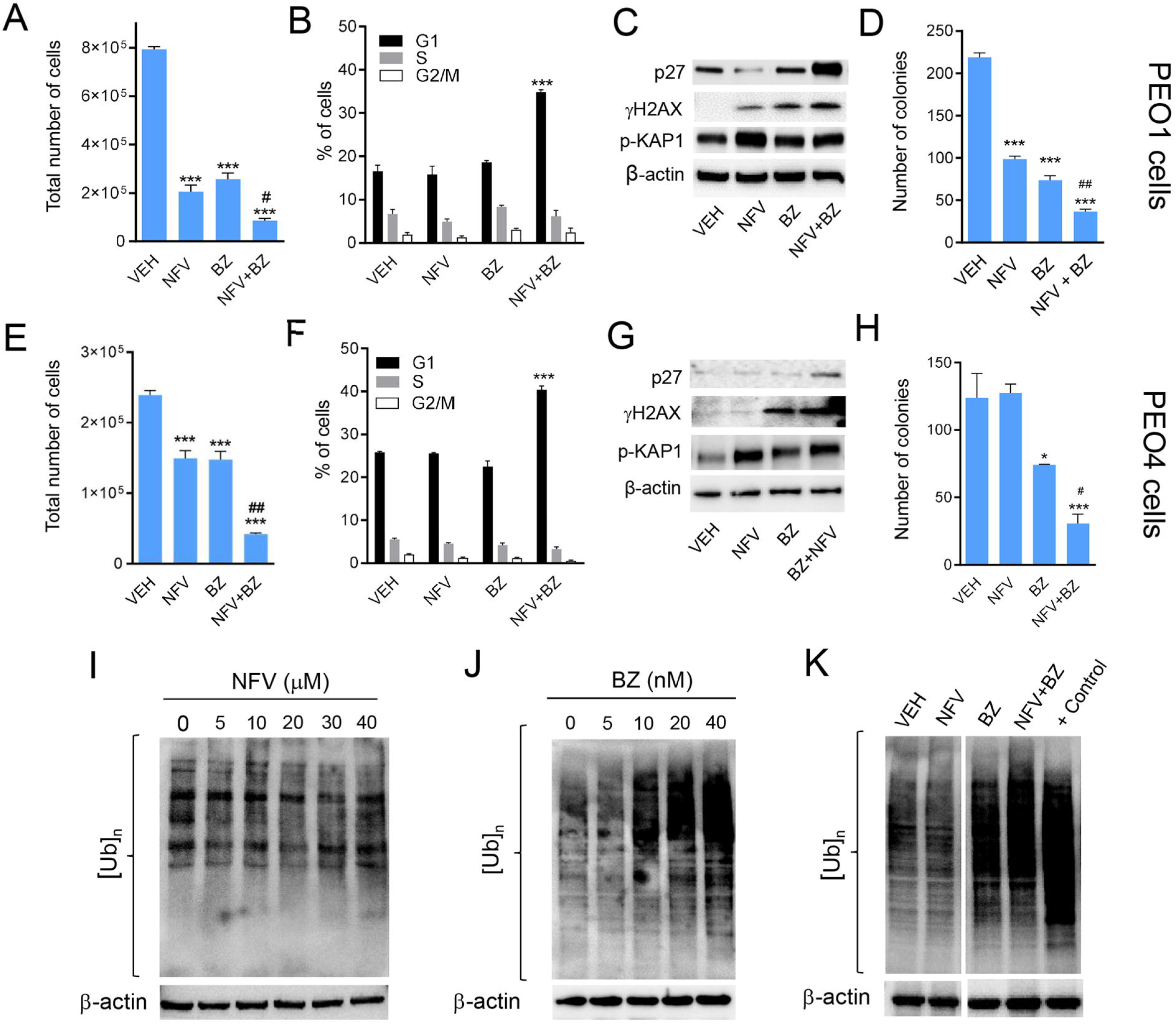
PEO1 and PEO4 were exposed to a combination of NFV and the proteasome inhibitor bortezomib (BZ) predicted to be synergistic in the drug interaction studies. Thus, PEO1 were exposed to 10 μM NFV, 10 nM BZ, or a combination of 10 μM NFV and 10 nM BZ. PEO4 instead received 5 μM NFV, 5 nM BZ, or the combination of 5 μM NFV and 5 nM BZ. In both cell lines, treatments lasted 72 h. Total number of PEO1 (A) or PEO4 (E) cells were assessed using a viability cell count reagent as described in materials and methods. In the same experiment, a fraction of cells obtained at the end of the experiment were stained with a solution of PI and subjected to cell cycle distribution analysis (B, PEO1 cells; F, PEO4 cells). Furthermore, 1000 live cells were collected at the end of the experiment and subjected to a clonogenic survival assay in drug-free media for 21 days (D, PEO1 cells; H, PEO4 cells). In a parallel experiment, total proteins were isolated and subjected to western blotting analysis for the expression of the p27^kip1^ cell cycle inhibitor and the DNA damage markers γH2AX and (Ser824) p-KAP1 (C, PEO1 cells; G, PEO4 cells). In A, D, E and H, *P<0.05 and ***P<0.001 compared against VEH, and #P<0.05 when compared NFV+BZ versus NVF or BZ alone. In B and F, ***P<0.001 compared to VEH. In separate experiments, PEO1 cells were exposed to increasing concentrations of NFV (I) or BZ (J) for 72 h and total proteins were isolated for western blot analysis of poly-ubiquitinated proteins ([Ub]n). In K, [Ub]n was measured in PEO1 cells exposed to either vehicle, 10 μM NFV, 10 nM BZ, or the combination of 10 μM NFV and 10 nM BZ for 72 h. A positive (+) control denoting high [Ub]n was generated from PEO1 ovarian cancer cells exposed to 20 nM BZ for 72. All experiments were repeated at least twice with a similar outcome.

We then subjected the cells that remained alive after 72 h of treatment with NFV, BZ, or the combination NFV/BZ—shown in Fig. 6A and E—to a clonogenic survival assay in the presence of drug-free media. We found that despite being alive after exposure to NFV and/or BZ, the cells were devoid of reproductive capacity as manifested by a reduction in their clonogenic survival, an effect manifested maximally when NFV was combined with BZ, when compared to the results of the drugs studied separately (Fig. 6D and H).

To determine whether the potentiation of the toxicity with the combination NFV/BZ was consequence of an enhanced inhibition of the proteasome by the drug combination compared with BZ alone, we measured the accumulation of poly-ubiquitinated proteins as a surrogate marker of proteasome inhibition. We observed that increasing concentrations of NFV did not change the rate of poly-ubiquitination (Fig. 6I). This lack of effect of NFV on the proteasome was also confirmed in PEO4, PEO14, and PEO23 cells (data not shown). As expected, we observed that BZ was able to increase protein poly-ubiquitination in a concentration-dependent manner (Fig. 6J). When we combined NFV and BZ at the concentrations that cause synergistic effects in terms of growth inhibition, decreased clonogenic survival, and increased levels of p27^kip1^, we confirmed that BZ alone but not NFV alone increased poly-ubiquitination. Most importantly, we observed that BZ-induced increase in poly-ubiquitinated proteins was not increased further by the presence of NFV (Fig. 6K). In summary, these results suggest that the potentiated toxicity between BZ and NVF is not consequence of furthering proteasome inhibition, yet it is associated with a potentiation in cell cycle arrest and reduced long-term clonogenic capacity likely consequence of unresolved DNA damage.

## Discussion

HGSOC is the most prevalent subtype of ovarian cancer, demonstrating heterogeneous phenotype and leading to the inevitable emergence of platinum resistance, which warrants the development of novel therapeutic avenues in the treatment protocol of this disease [1]. In this study, we explored the anticancer drug-repurposing potential of the HIV-PI compound nelfinavir (NFV), which has been in use to treat AIDS for over 20 years, demonstrating good tolerability as an anti-infective agent. Although NFV was demonstrated to be efficacious against multiple cancers [11], its role against the most deadly histopathological subtype of ovarian cancer, HGSOC, has been underexplored.

One of the challenges hindering the preclinical testing of novel treatments against HGSOC has been the lack of genomic similarity with the actual disease in the most-cited HGSOC cell lines [50]. One significant aspect of this study is the usage of patient-derived cell lines established longitudinally at different stages of disease progression and platinum sensitivities. These cell lines demonstrated differential morphological and genetic patterns *in vitro* [19, 20], and matched the genomic landscape of HGSOC [51]. We further describe that the cell lines recapitulated *in vitro* the status of the intrinsic cisplatin-sensitivity of the original patients (Table 1), providing clinical relevance to our study. All results reported using cells representing HGSOC should be considered in the context of heterogeneity of tumor genetics [52]. It is clear that this disease has clonal evolution beginning at its site of origin, which most evidence points to the serous tubal intraepithelial carcinoma (STIC) lesions that display genetic heterogeneity providing a platform for HGSOC evolution. STIC lesions were found closely associated with invasive HGSOC, with both lesions containing different *TP53* mutations, suggesting that the primary HGSOC may be clonally independent of the STIC [53]. The same paradigm arises when discussing clonal evolution between the primary lesion and different metastatic growths within the peritoneal cavity [52]. It is highly possible that there are different clones evolving during disease progression (i.e. parallel progression), and, in response to chemotherapy, the fittest clone that survives within the peritoneal cavity of recurrent patients may be selected. For the cell lines we used in this study, it is highly possible that PEO4 (or PEO23) cells obtained when the patients developed platinum resistance were already present in the initial disease when it still was platinum sensitive when the PEO1 (or PEO14) cells constituted the main clones. Likely, platinum drugs selected PEO4 or PEO23 cells from a pool of predominantly PEO1 or PEO14 cells, depending on the patient represented in our work. Thus, selecting HGSOC cell lines representing the different stages of disease evolution is of utmost relevance in order to develop the most efficient targeted therapies against the relevant clones dominating each disease stage [52, 53]. We report that NFV elicited effective cytotoxicity in all patient-matched HGSOC cell lines irrespective of their differential cisplatin sensitivity, evidenced by the concentration-dependent reduction in the total number of cells and percent viability, with the concomitant increase in hypodiploid-DNA content. Following short-term treatment with NFV for 72 h, the cells that remained alive were subjected to clonogenic survival assay in drug-free media to assess whether the cells recovered from the NFV-inflicted cytotoxicity. A concentration-dependent reduction in the number of clones suggested that the cells had sustained irreparable damage caused by NFV. This chronic toxicity may be explained by the induction, by NFV, of concentration- and time-dependent DNA damage as reflected by the increased phosphorylation of the DNA damage markers H2AX (a.k.a. γH2AX) and KAP1 [54, 55].

NFV demonstrated cytotoxicity in HGSOC cells originated from different patients, with different disease progression and cisplatin-sensitivity. Our results are in agreement with those of Brüning and colleagues showing NFV toxicity against ovarian cancer cells of different sensitivities to carboplatin [18] We therefore hypothesized that similar mechanistic pathways could be responsible for the generalized cytotoxic effects. Indeed, we observed a concentration-dependent increase in the expression of ER stress sensor GRP78, ER-stress related apoptosis mediator CHOP, and cell cycle inhibitor p27^kip1^ associated with inhibitory phosphorylation on Cdk2, suggesting that ER stress and cell cycle arrest are mechanisms activated in NFV-treated HGSOC cells, which contribute to the generalized cytotoxicity. Previously, Jiang *et al.* reported NFV-mediated upregulation of cell cycle inhibitor p27^kip1^ in melanoma cells, which accompanied reduced Cdk2 activity due to reduced Cdc25A phosphatase [17]. Furthermore, activation of the UPR upon ER stress has been associated with NFV mediated cytotoxicity against multiple cancers, such as lung cancer, glioblastoma, multiple myeloma, and breast cancer to mention some [12,15,56,57].

The UPR represents a series of signaling transduction events that ameliorate the accumulation of unfolded/misfolded proteins in the ER. It can have either a pro-survival or a proapoptotic role in the cells, depending on the intensity or the length of the stress [32,35,58–62]. Cancer cells have been reported to exploit ER stress for survival within unfavourable conditions, such as nutrient shortage, hypoxia, acidosis, and energy deprivation. As such, two pharmacological approaches can be used to take advantage of the chronically enhanced ER stress in cancer cells, either by shutting down the pro-survival mode of the UPR, or by tilting the cellular environment toward its proapoptotic phase [32, 36]. Our study demonstrated a concentration-dependent and temporal proapoptotic shift of the UPR in HGSOC cells in response to NFV treatment, which was evident from the enhanced expression of ER stress-related apoptosis mediators ATF4 and CHOP, accompanied with the increase of the Bax/Bcl2 ratio and a concomitant cleavage of executioner caspase-7. These results corroborate previous findings in non-small cell lung cancer and multiple myeloma cells, demonstrating the activation of the ATF4-CHOP pathway, and resulting in a proapoptotic shift of ER stress in response to NFV; the authors additionally reported a reduction of NFV-induced cell death during siRNA-mediated inhibition of CHOP, underpinning a key role of CHOP in the apoptotic process [63]. Pharmacological aggravation of constitutive ER stress by NFV in cancer cells has also been utilized as a chemosensitizing strategy against doxorubicin-resistant breast cancer and castration-resistant prostate cancer [14, 64].

A recent study shows high expression of GRP78, PERK, and ATF6 in patient’s tumors, which correlated with poor patient survival in HGSOC [65]. Elevated basal expression of ER stress-related proteins in ovarian cancer suggests the existence of a possible therapeutic window whereby further pharmacological aggravation of ER stress may induce apoptosis in ovarian cancer cells without triggering similar outcomes in normal cells. We observed that NFV might achieve such a goal as it activates all three arms of the UPR: IRE1α-XBP1, PERK-ATF4-CHOP, and ATF6 in HGSOC cells, to a comparable level as that caused by the classical ER stressor tunicamycin. Previously, it was reported an accumulation of ATF6 in prostate cancer cells due to NFV-mediated inhibition of site-2 protease (S2P) enzyme, which interrupted the regulated intramembrane proteolysis (RIP) of ATF6 in the Golgi apparatus for the release of the active soluble form [66]. Our result demonstrating NFV-associated increase in soluble ATF6 in HGSOC cells excludes the Golgi-resided S2P enzyme as a likely target of NFV in the HGSOC cells.

In this study, we reported a concentration- and time-dependent inhibition of protein synthesis by NFV, which was further abrogated by the presence of the protein synthesis inhibitor CHX. These results support the validity of the non-radioactive method for assessing mRNA translation that we used in this study and termed puromycin incorporation assay. Global protein synthesis inhibition was clearly the long-term outcome of NFV treatment in HGSOC cells; such effect, however, did not take place until after 4 h of NFV treatment. This was associated with a transient dephosphorylation of eIF2α and the cleavage of XBP mRNA, providing proof-of-principle that the initial sustained protein synthesis in the presence of NFV is, at least in part, responsible for triggering the UPR in HGSOC cells. We observed similar results in OV2008 ovarian cancer cells treated with the anti-progestin/anti-glucocorticoid agent mifepristone, which killed the cells because of an increase in ER stress that was associated with a short-term spike in protein synthesis that preceded the global abrogation of mRNA translation that concurred with the dying of the cells [26].

Other studies have shown that NFV can increase autophagy [12, 67]. In ovarian cancer cells, we have shown previously with the non-canonical ER stressor mifepristone that it caused ER stress-mediated toxicity by increasing autophagic flux and synergized with the lysosome inhibitor chloroquine in killing the cells [26]. In the case of NFV, while we observed an increase of LC3II in NFV-treated HGSOC cells of differential platinum sensitivities, suggesting an increase of the level of autophagosomes, we did not observe a further enhancement in the level of LC3II during co-treatment with the lysosomal inhibitor bafilomycin A. This signifies that NFV does not affect autophagic flux, but that the number of autophagosomes actually increases because of inhibition of lysosomal function.

We further report the reduction of survival and proliferation signals marked by the decline in p-AKT and p-ERK in HGSOC cells with high or low sensitivity to cisplatin, upon treatment with NFV. Downregulation of AKT is a well-known effect of NFV and has been associated with impaired glucose metabolism, insulin resistance, and lipodystrophy during chronic treatment, which are reversible upon discontinuation of the therapy [9]. This is relevant for a therapeutic standpoint, as amplified expression of components of the PI3K-AKT pathway has been correlated with reduced overall survival of HGSOC patients [68]. In patients with advanced HGSOC, amplification of the PI3KCA and AKT2 has been observed in 12% and 10% of samples, respectively [69] while reduced expression or loss of PTEN has been correlated with advanced staging in HGSOC samples [70, 71]. The inhibition of p-AKT by NFV was reported in other cancers, such as breast cancer [72, 73], multiple myeloma [15, 74], acute myeloid leukemia [75], pediatric refractory leukemia [76], diffuse B-cell lymphoma [77], prostate cancer [78], and non-small cell lung carcinoma [79]. The reduction in p-AKT by NFV has been also proposed as a radiosensitizing strategy in glioblastoma, bladder, lung, and head and neck cancers [48,80–82]. It is interesting to note that the reduction in p-AKT in peripheral blood mononuclear cells (PBMCs) was proposed as a surrogate biomarker to assess the pharmacological efficacy in targeting AKT signaling by NFV [83]. Also, p-AKT was decreased when NFV was combined with doxorubicin in doxorubicin-resistant chronic myeloid leukemia cells [84] and with the proteasome inhibitor BZ in multiple myeloma cells [74].

In terms of p-ERK inhibition, our data are commensurate with previous reports where reduction of ERK phosphorylation was observed in response to NFV in multiple myeloma [15, 85], HER2-positive and –negative breast cancer cells [13], medullary thyroid cancer [86], and adenoid cystic carcinoma [87].

Another significant finding in this study is the increase, upon NFV treatment, in the phosphorylation of H2AX (γH2AX)–a marker of DNA double strand breaks—in PEO1 and PEO4 cells having different sensitivities to cisplatin. Furthermore, we confirmed these results via the NFV-induced increase in the ATM-dependent phosphorylation of KAP1, which decondenses DNA upon DNA damage [55]. Our results concur with those obtained by Jensen and colleagues who demonstrated, in thyroid cancer cells, that similar doses of NFV than the ones used in our study induced the expression of γH2AX and p53BP1 indicating DNA damage [41].

HGSOC cells frequently present with TP53 mutations (97%) and a defect in the homologous recombination (HR) DNA repair mechanism (50%) primarily due to germline or somatic mutation of BRCA1/2 [88]. A deficient HR mechanism prevents error-free repair of DNA double strand breaks induced by platinum adducts, thus confers sensitivity of cancer cells to platinating agents. PEO1 cells carry a germline inactivating mutation to BRCA2 and are sensitive to platinating agents, whereas PEO4 that were obtained when the patient was resistant to platinum showed a functional restoration to the BRCA2 due to a secondary mutation [20, 89]. Deficiency in HR forces ovarian cancer cells to be over-reliant on the base-excision repair mechanism by poly (ADP-ribose) polymerase (PARP) classically utilized to repair single-strand DNA breaks. As such, targeting PARP has been a desirable pharmacologic approach to induce synthetic lethality in ovarian cancer cells [90]. It has also been implicated that restoration of BRCA2 confers cross-resistance to PARP inhibitors in parallel to reduced cisplatin sensitivity [91]. Since NFV was able to elicit enhanced γH2AX and p-KAP1 in HGSOC cells independent of their BRCA status and sensitivity to cisplatin, a different mechanism of DNA damage may be involved whereby the cells may not rely on the HR pathway to repair their DNA.

Finally, we report that NFV can potentiate the effects of proteasome inhibitor BZ by affecting the cell cycle in combination with sustained DNA damage. Inhibition of the proteasomes leads to the accumulation of ubiquitinated proteins, which in turn can increase the protein load in the ER and push the effect of an ER stressor to lethality. Based on this rationale, we demonstrated that the non-canonical ER stressor mifepristone potentiates the effect of BZ in ovarian cancer cells by significantly inhibiting the activity of the proteasome leading to cell death [26]. In the present study using PEO1 and PEO4 HGSOC cells, however, NFV, despite causing ER stress similarly to mifepristone, it did not inhibit the proteasome beyond the inhibition caused by BZ alone. This is not surprising as previous reports suggested that the effect of NFV on the proteasome may be cancer and cell-type specific; for instance, NFV did not demonstrate an inhibitory effect on the proteasome in cervical cancer cells [92]. What we found when combining NFV and BZ in the current HGSOC cells, was an increase in the expression of the cell cycle inhibitor p27^kip1^, which was higher than the expression induced by each drug individually; this was associated with a potentiation among the drugs in causing cell cycle arrest at the G1 phase of the cell cycle. In the long-term, likely because of sustained DNA damage, the combination NFV/BZ cause reduced clonogenic survival, suggesting a potentiation among the drugs irreversibly abrogating the reproductive capacity of the cells. The potentiation of toxicities between NFV and BZ was also reported in non-small lung cancer cells and multiple myeloma cells [63], as well as in leukemia cells [75]. Other combination reported to lead to a potentiation of effects with NFV is that of the anti-diabetic drug metformin, which together with NFV exhibited greater inhibition of growth of cervical cancer cells *in vitro* and *in vivo* in nude mice than either NFV or metformin alone [93]. Platinum resistance is one of the primary reasons that has interrupted the improvement of patient survival in advanced stage HGSOC. Hence, a drug targeting the platinum-resistant phenotype is highly desirable in research for newer therapies for this disease. Our data demonstrate that NFV induces toxicity toward HGSOC cells of a wide-range of platinum sensitivities possibly through a DNA-damaging mechanism, which is likely different to that caused by cisplatin as we did not find cross-resistance among the drugs. One challenge in introducing NFV in the clinical practice for treating HGSOC would be maintaining the desired plasma concentration that achieve anticancer effects. The anti-infective dosing of 1250 mg twice daily yielded a wide range of variability in the peak plasma concentration (4.4 - 11.3 mg/L), likely due to genomic polymorphism in the metabolizing enzyme CYP2C19 [94]. Other authors have reported the concentration reached in circulation by NFV of being in the 7.7 to 20 μM range [41], which is within the range of concentrations we used in the current study. Chemical hybridization and combination with other drugs may be used to keep optimal plasma concentration of NFV to achieve greater anticancer efficacy. For instance, recently, nitric oxide hybridization of HIV-PIs has been promoted as an alternative strategy to improve pharmacokinetics and anticancer efficacy [95].

## Conclusions

In summary, in this study we demonstrated multipronged mechanisms whereby NFV targets HGSOC cells independent of their platinum sensitivities (Fig. 7). As an oral anti-infective drug with a well-documented history of tolerable side effects, the observed anticancer effects of NFV suggest its potential repurposing benefit against HGSOC as an additional adjuvant chemotherapeutic agent.

**Figure 7.**
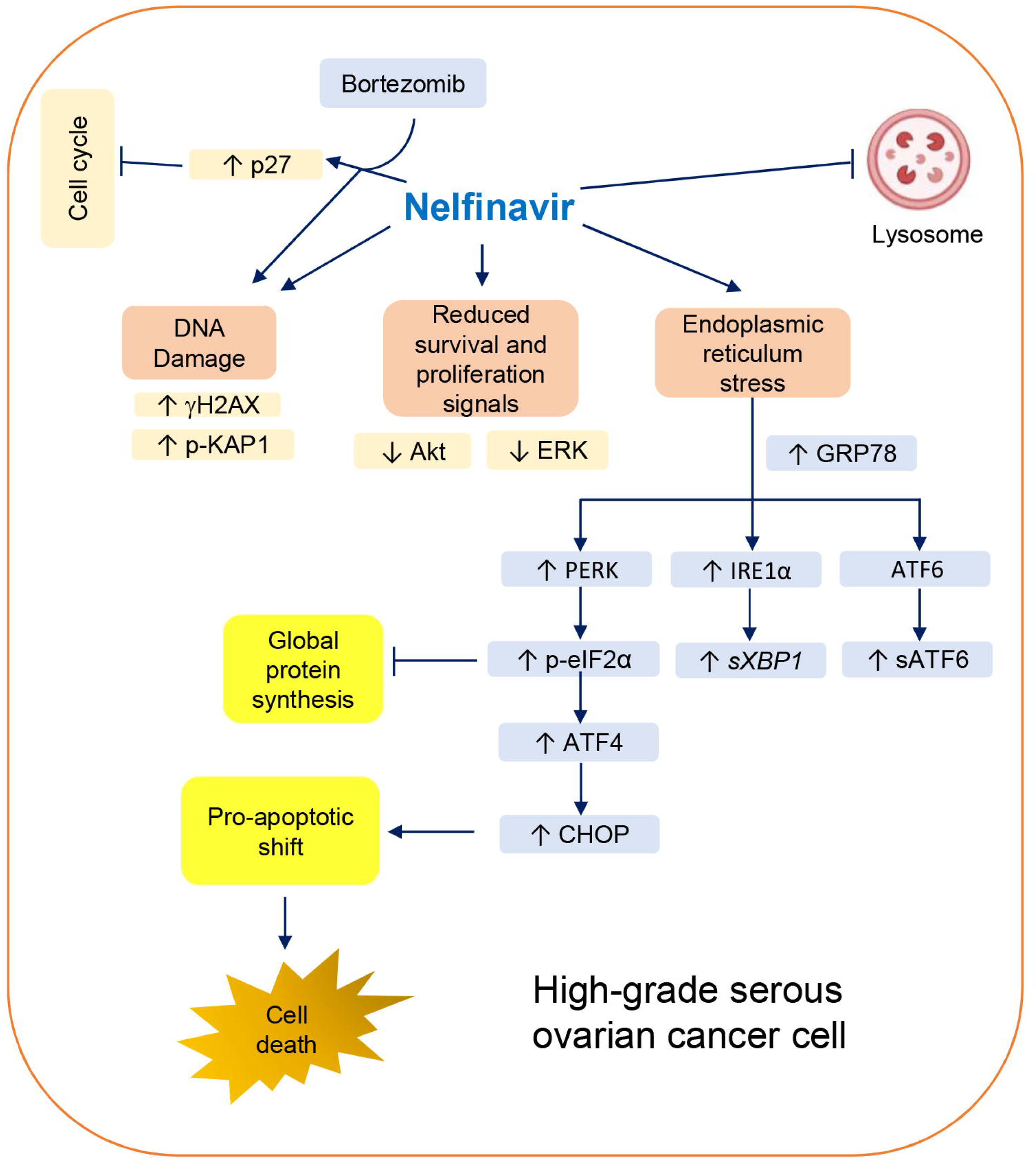
Schematic representation of the proposed mechanism of NFV-mediated cytotoxicity in HGSOC cells regardless of platinum sensitivity. NFV triggers DNA damage, reduces survival and proliferation signaled by AKT and ERK, and activates the three arms of the UPR: 1) IRE1α- XBP1, 2) PERK-eIF2α-ATF4-CHOP, and 3) ATF6 cleavage. Phospho-eIF2α leads to inhibition of global protein synthesis, and, downstream of it, ATF4-CHOP-mediated proapoptotic shift triggers the cleavage of executioner caspase-7 to elicit cell death. Additionally, NFV impairs the autophagic clearance, likely via lysosome inhibition. Finally, NFV potentiates the cytotoxicity of proteasome inhibitor BZ by increasing the level of cell cycle inhibitor p27^kip1^, while reducing clonogenic survival (not shown) denoting a long-lasting toxicity likely consequence of sustained DNA damage.

## Acknowledgments

This research was supported by a grant from the Rivkin Center for Ovarian Cancer Research (Seattle, WA, USA), and funds from the Department of Pathology, Faculty of Medicine and Health Sciences, McGill University. We are indebted to Millennium Pharmaceuticals for providing bortezomib (Velcade®) for preclinical research purposes.

## Supplementary Materials

**Table S1:** Source and dilutions of antibodies used; **Table S2:** IC_50s_ of clonogenic inhibition by NFV of the cell lines studied in this work: **Figure S1:** Induction of p27^kip1^ and reduction in phosphorylation of Cdk2 upon NFV treatment; **Figure S2:** Effect of NFV on the accumulation of LC3II in PEO14 and PEO24 cells; **Figure S3:** Cytotoxicity of NFV towards HGSOC cells in the presence or absence of small molecule salubrinal;

## Notes

### Competing Interest Statement

The authors have declared no competing interest.

### Summary of Updates

This version of the manuscript includes some new data, for instance the confirmation of DNA damage induced by nelfinavir upon Ser824 phosphorylation on KAP1. It also contains data on the Thr308 phosphorylation of AKT upon nelfinavir treatment. There is also calculation of IC50s of the clonogenic survival of high-grade serous ovarian cancer cells in response to nelfinavir. Also, we demonstrate that the toxicity of nelfinavir leads to increase of p27 in association with a dephosphorylation of Cdk2 on Thr160. Finally, we include data demonstrating that the small molecule ER stress inhibitor salubrinal, is able to abrogate the cytotoxicity induced by nelfinavir. More articles are also included in the discussion as well as an entire new paragraph on the clonal heterogeneity of high-grade serous ovarian cancer.

