## Supplementary Figures for "Nelfinavir induces cytotoxicity towards high-grade serous ovarian cancer cells, involving induction of the unfolded protein response, modulation of protein synthesis, DNA damage, lysosomal impairment, and potentiation of toxicity caused by proteasome inhibition"

**Table S1.** Source and dilutions of antibodies utilized in this work.

| Antibody | Clone or catalogue | Company | Concentration |
|---|---|---|---|
| PERK | 3192S | Cell Signaling Technology | 1:1000 |
| GRP78 | 3177S | Cell Signaling Technology | 1:1000 |
| IRE1α | 3294S | Cell Signaling Technology | 1:1000 |
| p-eIF1α | 3398S | Cell Signaling Technology | 1:1000 |
| eIF2α | 9722S | Cell Signaling Technology | 1:1000 |
| ATF4 | 11815S | Cell Signaling Technology | 1:1000 |
| CHOP | 5554S | Cell Signaling Technology | 1:1000 |
| ATF6 | NBP1-40256 | Novus Biologicals | 1:1000 |
| Puromycin | MABE343 | EMD Millipore Corporation | 1:20000 |
| p-Akt (Ser473) | 4058L | Cell Signaling Technology | 1:1000 |
| p-Akt (Thr308) | 4056L | Cell Signaling Technology | 1:1000 |
| Akt | 2920S | Cell Signaling Technology | 1:1000 |
| p-ERK | 9106L | Cell Signaling Technology | 1:1000 |
| ERK | 9102L | Cell signaling Technology | 1:1000 |
| p27 | 610242 | BD Biosciences | 1:1000 |
| γH2AX | 05-636 | EMD Millipore Corporation | 1:1000 |
| p-KAP1 (Ser824) | NB100-2350 | Novus Biologicals | 1:1000 |
| LC3B | 3868S | Cell Signaling Technology | 1:1000 |
| Cdk2 | sc-6248 | Santa Cruz Biotechnology | 1:500 |
| p-Cdk2 (Thr160) | 2561S | Cell Signaling Technology | 1:1000 |
| Ubiquitin | 3933S | Cell Signaling Technology | 1:1000 |
| β-Actin | A5441 | Sigma Life Sciences | 1:10000 |
| Caspase-7 | 12827S | Cell Signaling Technology | 1:1000 |
| Bax | 5023S | Cell Signaling Technology | 1:1000 |
| Bcl-2 | 15071S | Cell Signaling Technology | 1:1000 |
| Anti-mouse | 170-6516 | BioRad Laboratories Inc. | 1:8000 |
| Anti-rabbit | 170-6515 | BioRad Laboratories Inc. | 1:10000 |

**Table S2.** Concentration of nelfinavir (NFV) needed to achieve 50% reduction in clonogenic survival ($IC_{50}$) of the HGSOC cell lines studied.

| Cell Line | $IC_{50}$ (μM) |
|---|---|
| PEO1 | 10.05 ± 0.75 |
| PEO4 | 12.40 ± 2.60 |
| PEO6 | 11.40 ± 0.40 |
| PEO14 | 16.07 ± 4.90 |
| PEO23 | 11.14 ± 0.71 |

**Figure S1**. Inhibition of Thr160 phosphorylation of Cdk2 (p-Cdk2) by nelfinavir (NFV) correlates with increased levels of Cdk inhibitor $p27^{kip1}$. PEO1 cells were incubated with 20 μM NFV for the depicted times. At the end of the experiment, total protein extracts were obtained, and western blotting assessed the expression of p-Cdk2, Cdk2, $p27^{kip1}$, and β-actin.

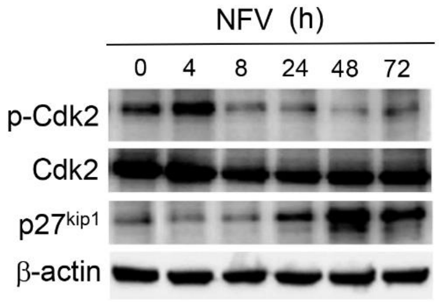


**Figure S2.** In PEO14 and PEO23 cells treated with various concentrations of nelfinavir (NFV) for 72 h; the autophagosome-related protein LC3II increases in response to NFV in a concentration-dependent manner.

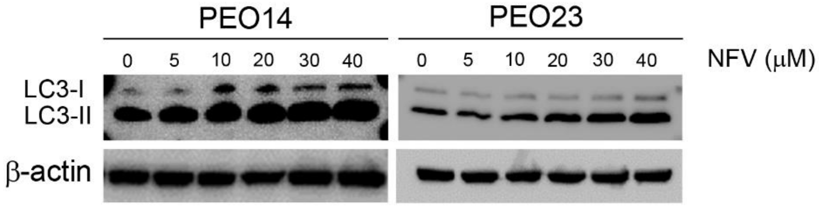

**Figure S3.** The cytotoxicity of nelfinavir (NFV) towards PEO1 ovarian cancer cells is significantly prevented by the small molecule salubrinal. PEO1 cells were treated vehicle (VEH) or with 20 μM NFV, with or without concurrent 50 μM salubrinal (SAL) for 72 h. At the end of the experiment, the cells were trypsinized and subjected to microcytometry analysis to attain their viability. *$p<0.05$ compared to NFV (One-way ANOVA followed by Tukey's Multiple Comparison Test).

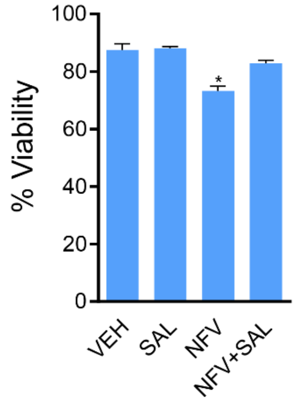
